# Cognitive regulation of multimodal food perception in the human brain

**DOI:** 10.1101/2020.04.23.057364

**Authors:** R. Janet, A. Fournel, M. Fouillen, E Derrington, M Bensafi, JC. Dreher

## Abstract

The ability to regulate appetite is essential to avoid food over-consumption. The desire for a particular food can be triggered by its odor before it is even seen. Using fMRI, we identify the neural systems modulated by cognitive regulation when experiencing appetizing food stimuli presented in both olfactory and visual modalities, while being hungry. Regulatory instruction modulated bids for food items and inhalation patterns. Distinct brain regions were observed for up and down appetite-regulation, respectively the dorsomedial prefrontal cortex (dmPFC) and dorsolateral PFC. Food valuation engaged the ventromedial PFC and bilateral striatum while the amygdala was modulated by individual food preferences, indexed by rank-ordered bids. Furthermore, we identified a neurobiological marker for up-regulating success: individuals with higher blood levels of ghrelin were better at exercising up-regulation, and engaged more the dmPFC. This characterizes the neural circuitry regulating food consumption and suggests potential hormonal and neurofunctional targets for preventing eating disorders.

## Introduction

Eating disorders such as binge eating disorder represent a public health challenge because they are associated with high comorbidity and serious health consequences (Hoek et al., 2016). The rising rates in obesity (Flegal et al., 2016; Gallus et al., 2015; Sturm et al., 2013) also emphasize the crucial need to understand the neurocomputational mechanisms underlying the regulation of food consumption, especially in a context of overexposure to food stimuli. Food consumption may be regulated through the implementation of strategies, known as cognitive regulation. These strategies use attention, language and executive control to modulate the value people attribute to features of visual stimuli (Adcock et al., 2006; Galsworthy-Francis and Allan, 2014; Yokum and Stice, 2013). Cognitive regulation thus serves as an important strategy by which the brain can control food craving.

Most of our knowledge about the neurobiological mechanisms underlying cognitive regulation of food stimuli is derived from fMRI studies using food images only (Hutcherson et al., 2012; Inui et al., 2004; Kober et al., 2010). These studies reported that presentation of visual food-cues engages brain regions associated with reward and valuation, the bilateral striatum and the ventromedial prefrontal cortex. Accumulated evidence indicate that the lateral prefrontal cortex (lPFC) also plays a key role in modulating food cue induced signals by the use of cognitive strategies (Hollmann et al., 2012; Hutcherson et al., 2012b; Kober et al., 2010; Siep et al., 2012a; Yokum and Stice, 2013 Schmidt et al., 2018). Early studies reported that the lPFC, supporting cognitive regulation, acts on value signals encoded in the ventromedial PFC (vmPFC) (Hare et al., 2009; Kober et al., 2010, Hutcherson et al., 2012b). However, a recent study found that the vmPFC activity may be unsensitive to regulatory goals during cognitive regulation (Tusche and Hutcherson, 2018), suggesting that cognitive regulation acts upstream of the integrated value signal by modulating specific attributes value (Inzlicht et al., 2016).

Yet, little is known about the neurobiological mechanisms underlying appetite regulation of food stimuli presented in a more realistic and ecological fashion, such as when combining visual cues with food odors. Food odors are potent signals for triggering appetite. For example, the smell of a croissant wafting from a patisserie, can trigger a strong desire for this food in the absence of any visual cue. Parallels may exist between the neural mechanisms engaged in cognitive strategies used for emotion regulation and for the regulation of appetite (Buhle et al., 2014; Kober et al., 2010). Both types of regulatory mechanisms may engage common brain regions, such as the dorsolateral PFC (dlPFC) for down-regulation (Frank et al., 2014) and the anterior medial part of the PFC for up-regulation (Ochsner et al., 2004). We therefore hypothesized that the lateral PFC may play a role in down-regulating food craving whereas the medial part of the PFC may support up-regulation of appetite.

Furthermore, homeostatic peptide hormones such as ghrelin, produced in the gastrointestinal tract and leptin, produced in adipose cells and the small intestine, convey energy balance information to the brain that affect food intake. Ghrelin acts both on the homeostatic hypothalamic-brainstem circuits regulating energy balance and on systems involved in reward and motivation (Mason et al., 2013; Perello and Dickson, 2015). High levels of ghrelin, either due to ghrelin injection or fasting, increase motivation for food rewards and modulate the reward system (Abizaid et al., 2006; Han et al., 2018; Karra et al., 2013; Kroemer et al., 2013; Shirazi et al., 2013). However, it remains unknown whether ghrelin modulates the brain systems engaged in appetite up-regulation in humans.

Here, we used fMRI to investigate the neural processes involved in appetite regulation during successive presentation of food odor and image. Healthy hungry participants made real food purchase decisions under a Natural control condition and two cognitive regulation conditions after smelling a food odor paired with a corresponding food picture. We investigated: 1) whether cognitive regulation modulates sniffing and subjective preferences for food, as assessed by willingness to pay; 2) which brain areas support valuation of food cues presented sequentially in the olfactory and visual modalities; 3) whether distinct or common PFC areas support up- and down-regulation of salient food items; 4) whether brain regions involved in the valuation of food cues are functionally modulated by regulatory regions; 5) whether individual preferences for specific odors have a specific neural signature (by rank ordering each food category according to each participants’ preferences); 6) what are the contributions of ghrelin and leptin to appetite regulation under cognitive regulation.

## Results

### Behavior

Prior to scanning, participants rated their appetite on a continuous scale (ranging from “0”= not hungry at all, 50= moderately hungry; to “100”= never been so hungry). The food quantity participants would be willing to eat prior to scanning was also assessed on a similar continuous scale. Participants rated their appetite at 60.4% (SEM= 4.7) and their food quantity at 75.1 (SEM= 3.2). This procedure allowed us to ensure that participants felt subjectively hungry and were willing to eat a large quantity of food. We also found a positive correlation between participants’ appetite and their blood level of ghrelin (r=0.495, p=0.016) and a positive trend between blood level of ghrelin and the quantity of food participants were willing to eat (r=0.406, p =0.054).

We found that cognitive regulation had a significant effect on average bidding behavior (F _(2,46)_ = 240.951, p<0.001) (Figure 2A). *Post hoc* analysis revealed that, compared to the Natural condition (the no cognitive regulation condition) (mean (M) 1.52 ± (SEM) 0.011), participants bid significantly more under the Indulge (the positive cognitive regulation condition) ((M) 2.03 ± (SEM) 0.011; paired t(_24_) = 12.508 p<0.001) and less under the Distance condition (the negative cognitive regulation condition) ((M) 0.871 ± (SEM) 0.007; paired t(_24_) = 11.197 p<0.001) (paired t-test with Bonferroni correction). We also controlled for possible interactions between regulatory conditions and food odor category (Apricot, Pineapple, Milk Chocolate or Dark Chocolate). This analysis revealed no significant interaction between cognitive regulation and types of food odor categories (F _(3,72)_ = 0.934, p=0.469). This analysis also revealed that participants’ bids differed according to the food category (F _(3,72)_ = 118.079, p<0.001) (Figure 2B). Post-hoc tests revealed that participants bid less for Pineapple ((M) 1.32 ± (SEM) 0.016) compared to milk chocolate ((M) 1.32 ± (SEM) 0.016) (p < 0.005, paired t_(24)_ = −3.145) and compared to dark chocolate ((M) 1.32 ± (SEM) 0.016) (p < 0.05, paired t_(24)_ = −2.789).

**Figure 1:**
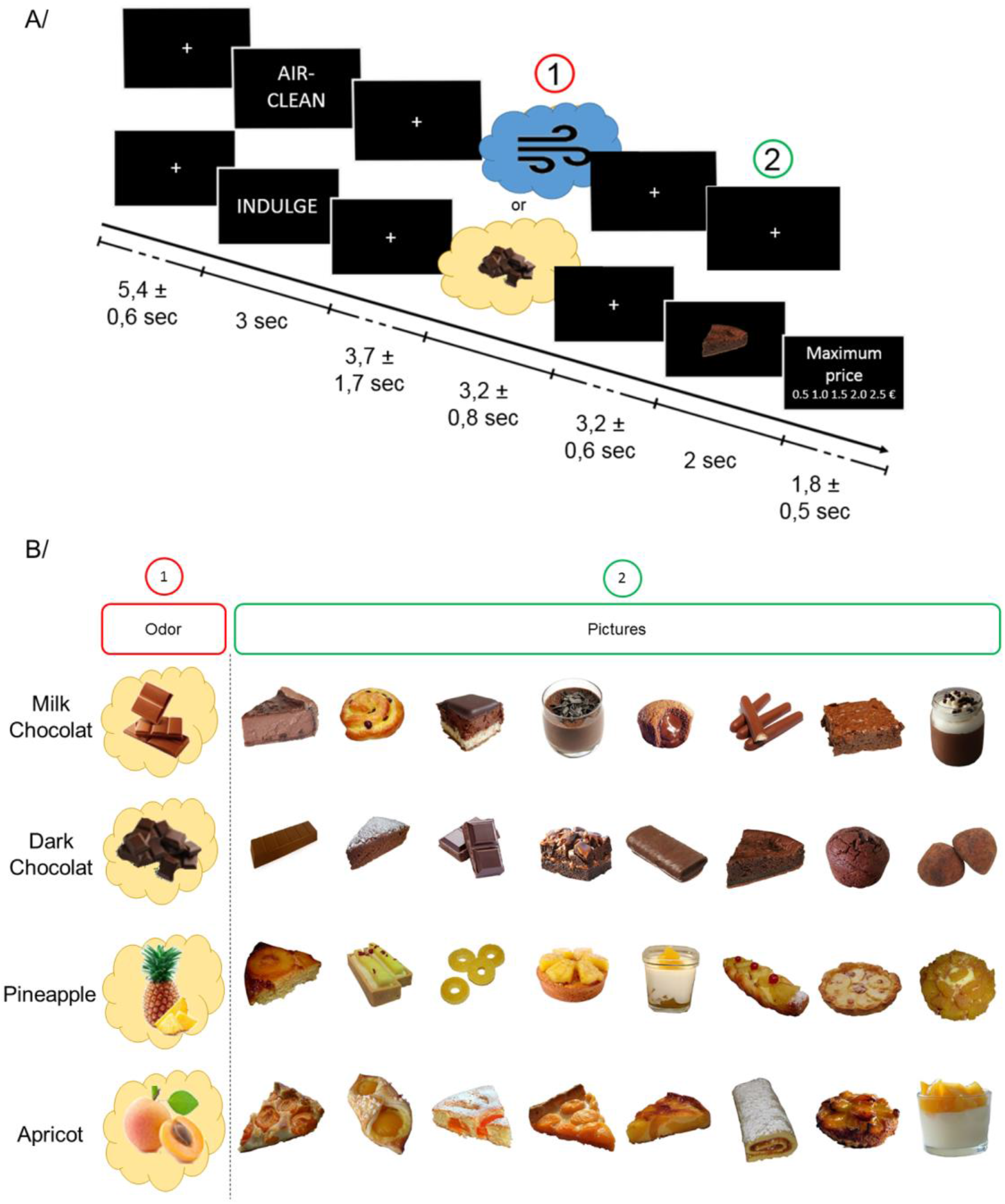
Experimental Design. (A) Schematic overview of one trial of the task. The task was composed of four steps. First, hungry participants were given instructions (Indulge, Natural or Distance) to regulate their craving towards food items. Second, they smelled one out of four odor categories (Apricot, Pineapple, Milk Chocolate or Dark Chocolate). Third, a picture of a food item associated to the odor was displayed (8 food pictures *per* odor category). Finally, participants were asked to rate how much they wanted to pay to get the food on a 5 points rating scale (from 0.5€ to 2.5€ with increment steps of 0.5€). (B) Overview of the food items presented. On each trial, a combination of one odor and one congruent picture (from the same food category) was presented.

**Figure 2:**
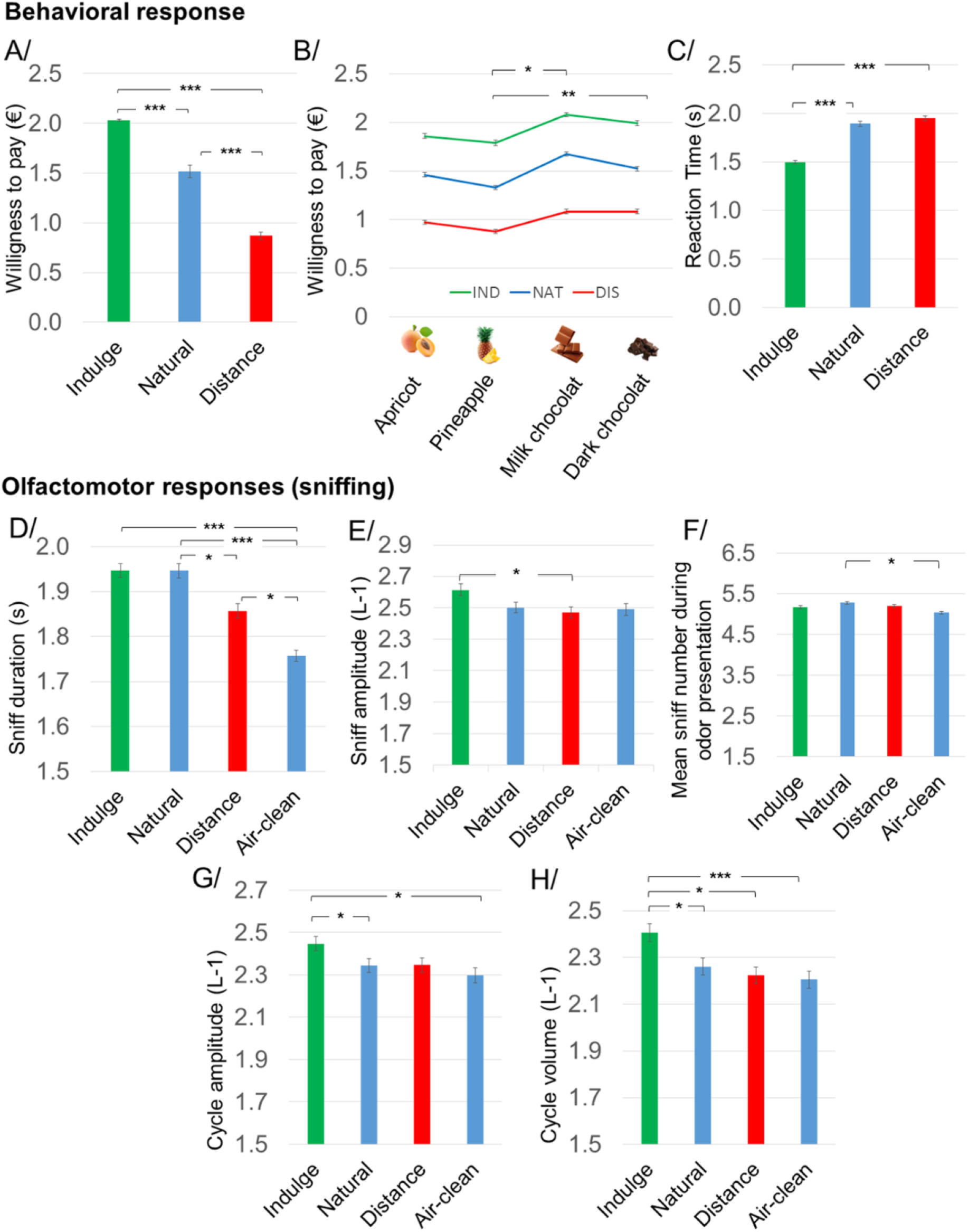
Behavioral and physiological influence of cognitive regulation. (A) Average bid across conditions of cognitive regulation. Bids decreased in the Distance condition compared to Natural, while bids increased in the Indulge condition. (B) Average bids across conditions and odor categories. Participants were less willing to pay for pineapple as compared to milk chocolate and dark chocolate. (C) Average Reaction Times. A decrease in RTs was observed in the Indulge compared to the Natural and Distance conditions. (D) Average duration of the first sniff. The duration of the first sniff was shorter in the Distance condition. (E) Average amplitude of the first sniff. The average amplitude of the first sniff was greater in the Indulge condition compared to the Distance condition. (F) Mean number of sniffs during odor presentation. Higher number of sniffs in the Natural compared to Air-Clean trials. (G) Average amplitude of sniffs across the entire period of sniffing. Amplitude was greater in the Indulge condition compared to Natural and Air-Clean conditions. (H) Average volume of sniffs across the entire period of sniffing. Volume was greater in the Indulge condition compared to Natural, Distance and Air-Clean conditions. Error bars show SEM. *** means p<0.001 ** means p<0.01 and * means p<0.05.

Reaction Times (RTs) also differed between conditions (F_(2,46)_ = 16.247, p<0.001) (Figure 2C). Participants’ bids were faster in the Indulge condition ((M) 1.50 ± (SEM) 0.015) compared to the Natural ((M) 1.90 ± (SEM) 0.024; paired t_(24)_ = 5.825 p<0.001) and Distance conditions ((M) 1.95 ± (SEM) 0.025) (paired t_(24)_ = 5.221 p<0.001).

### Olfactomotor responses (sniffing)

A significant effect of regulatory instructions was observed on the duration of the first sniff (Friedman test, χ^2^_(3)_ = 30.064, p = 0.001) (Figure 2D, Duration of first sniff). There was a significant decrease in the duration of sniffing in the Distance ((M) 1.86 ± (SEM) 0.015) compared to the Natural condition ((M) 1.94 ± (SEM) 0.016) (p =0.011, paired t_(24)_ = −2.92). Participants inhaled for a shorter period during the Air-clean ((M) 1.76 ± (SEM) 0.012) condition compared to the Distance, Indulge ((M) 1.95 ± (SEM) 0.016) and Natural conditions. No significant difference in sniffing duration was found between the Indulge and Natural conditions (paired t_(24)_, = 2.587, p = 0.096).

A significant effect of regulatory instructions was observed concerning the amplitude of the first sniff (F _(3,72)_, = 4.248, p = 0.008) (Figure 2E, Amplitude of first sniff). A *post-hoc* test with the Bonferroni correction showed a significant decrease in the amplitude of sniffing in the Distance ((M) 2.47 ± (SEM) 0.035) compared to the Indulge condition ((M) 2.61 ± (SEM) 0.033) (p = 0.025, paired t_(24)_ = −3.139). No significant effect of condition was observed on the volume of the first sniff (F _(3,72)_, = 2.587, p =0.06).

We next considered the entire sniffing period during odor presentation. We found a significant difference between conditions with respect to the total number of sniffing cycles (F _(3,72)_, = 3.962, p = 0.011) (Figure 2F, Mean sniff number). *Post-hoc* analysis revealed that participants had more sniffing cycles in the Natural condition compared to the Air-clean condition (p = 0.024, paired t_(24)_ = −2.65).

A significant effect of conditions was observed on the average sniffing amplitude during the total sniffing period (F _(3,92)_, = 6.068, p = 0.001) (Figure 2G, cycle amplitude). A *post-hoc* test with Bonferroni correction showed a significant increase in the average amplitude of sniffing in the Indulge ((M) 2.45 ± (SEM) 0.035) compared to the Natural conditions ((M) 2.34 ± (SEM) 0.035) (p = 0.037, paired t(_24_) = −2.448). The *Post-hoc* test also revealed a significant increase in the amplitude of sniffing in the Indulge compared to the Air-clean conditions ((M) 2.35 ± (SEM) 0.034) (p = 0.011, paired t(_24_) = −2.346). Finally, no significant difference concerning the sniffing amplitude was observed between the Indulge and Distance conditions.

A significant effect of conditions was observed on the average sniff volume during the total sniffing period (F _(3,72)_, = 7.863, p = 0.001) (Figure 2H, Mean sniff volume). A *post-hoc* test with the Bonferroni correction showed a significant increase in the mean volume of sniffing in the Indulge ((M) 2.41 ± (SEM) 0.037) compared to the Natural condition ((M) 2.26 ± (SEM) 0.036) (p = 0.028, paired t(_24_) = −2.448). They also revealed an increase in the average volume of sniffing in the Indulge compared to Distance conditions ((M) 2.22 ± (SEM) 0.036). Finally, the *post-hoc* test also revealed a significant increase in the average volume of sniffing in the Indulge condition compared to the Air-clean trials ((M) 2.20 ± (SEM) 0.036) (p = 0.004, paired t(_24_) = −2.346).

No significant effect of conditions was observed concerning the mean duration of sniffing during the entire period of odor presentation (F _(3,72)_, = 1.313, p = 0.277).

We also investigated potential differences between men and women in their capability to regulate their appetite. Using a Two-way ANOVA, we found no significant difference between sex concerning bid (F(_5,120_) = 4.229, p = 0.132) or RTs (F_(5,120)_ = 5.603 p = 0.099). When performing the same analysis on breathing parameters, we observed no significant differences for the volume (F_(7,168)_ = 0.014; p = 0.907), amplitude (F_(7,168)_ = 1.31; p = 0.282) and the duration of the first sniff (F_(7,168)_ = 4.42; p = 0.065). Despite the fact that men have a greater total lung capacity (LoMauro and Aliverti, 2018) we found no differences between men and women in breathing parameters.

#### fMRI results

##### Neurocomputational mechanisms of cognitive regulation of food

First, using the GLM1, we searched for brain regions engaged in cognitive regulation during the odor/image presentation. As shown in Figure 3, Table 1, when averaging over the period of odor/image presentation, BOLD response in the dorsomedial PFC was significantly higher in the Indulge compared to Natural condition (comparison Indulge > Natural) (x,y,z: −4, 58, 21; t = 5.43; p<0.05 Family-Wise Error (FWE) whole brain cluster corrected). To illustrate the response in this brain region, we extracted beta parameters in the three different conditions and plotted them (Bar Graphs). We also investigated the opposite contrast (Natural > Indulge). No brain region showed greater activity under Natural trials compared to Indulge trials.

**Figure 3:**
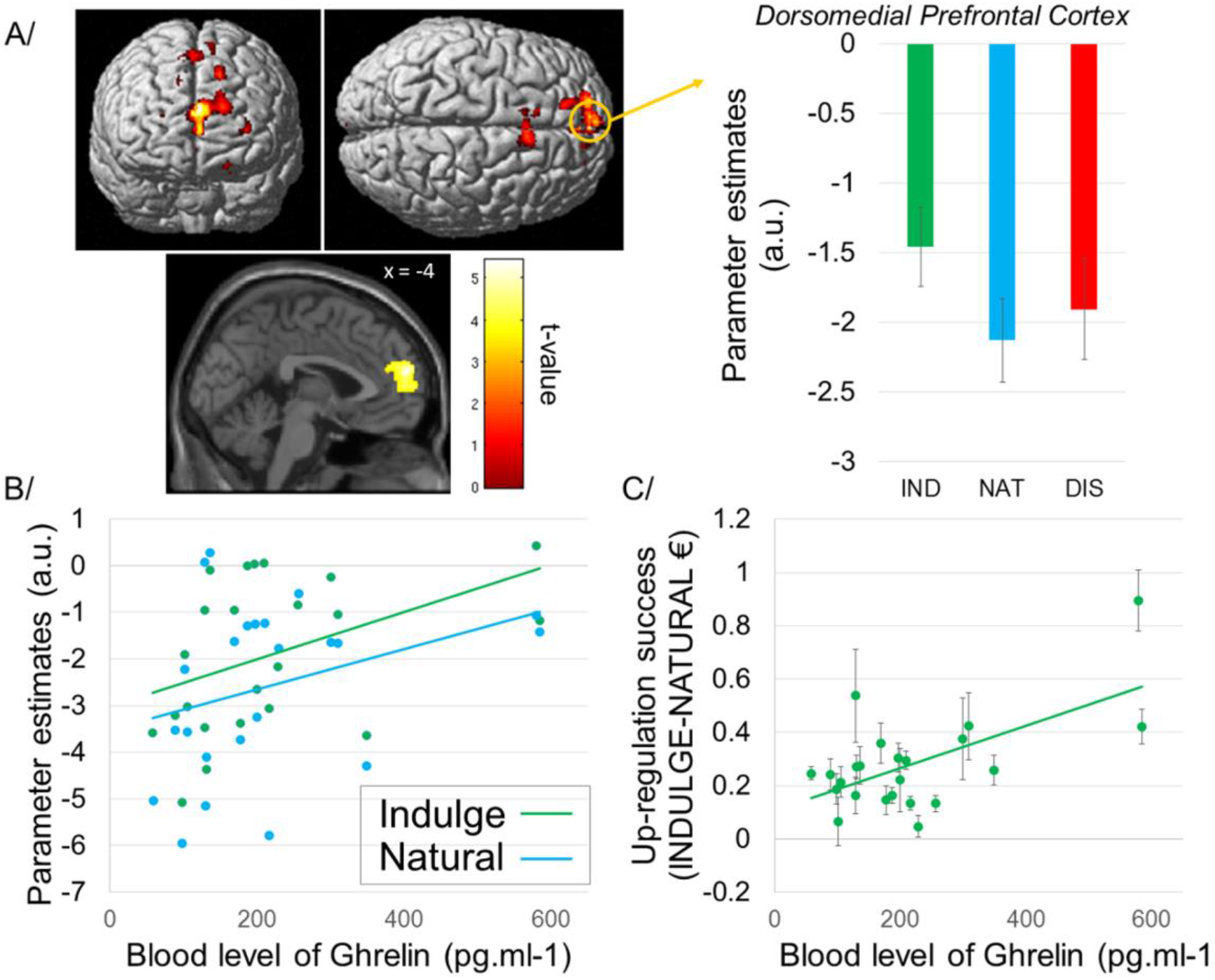
Up-regulation of appetite increases dmPFC activity and inter-individual differences linking up-regulation success and ghrelin levels. (A) Up-regulation during the Indulge condition increased activity in the dorsomedial PFC (x,y,z: −4, 58, 21), at a voxel-wise threshold of p<0.001 and cluster size > 40, corresponding to a whole brain FWE cluster corrected threshold of p<0.05. Error bars show SEM. (B) Positive correlation between blood level of ghrelin and parameter estimates from the Indulge and Natural conditions. Correlation between beta in the Natural condition and ghrelin level r=0.469 (p=0.024) and between beta from the Indulge condition and ghrelin level r=0.511 (p=0.013). (C) Positive correlation between up-regulation success and ghrelin level (r=0.373, p<0.05). Participants showing higher levels of ghrelin were also those showing the higher up regulatory success, as they were willing to pay even more during the Indulge condition.

**Table 1:** BOLD changes induced by cognitive regulation during processing of food items. **Clusters are reported at p < 0.05, family-wise error (FWE) cluster-level corrected for multiple comparisons (with an initial cluster-forming threshold of p < 0.001 and an extent k = 40 voxels).

| Effect of cognitive regulation during<br>food perception | MNI peak cluster coordinates |  |  |  |  |
| --- | --- | --- | --- | --- | --- |
|  | x | y | z | k | Z score |
| Indulge > Natural |  |  |  |  |  |
| Right Superior Medial frontal ** | -4 | 58 | 21 | 1505 | 4.31 |
| Left Superior Medial frontal ** | -32 | 38 | 12 | 317 | 4.01 |
| pre-SMA ** | 8 | 16 | 54 | 271 | 3.78 |
| Anterior Cingulate | 0 | 32 | -3 | 75 | 3.64 |
| Left Superior Medial frontal | -16 | 45 | 45 | 131 | 3.59 |
| Indulge < Natural |  |  |  |  |  |
| No brain region |  |  |  |  |  |
| Distance > Natural |  |  |  |  |  |
| Left Superior Lateral frontal ** | -24 | 52 | 22 | 6343 | 4.97 |
| Dorsolateral Prefrontal cortex ** | 44 | 16 | 46 | 561 | 4.27 |
| Right Angular ** | 56 | -54 | 27 | 722 | 4.22 |
| Left Angular ** | -46 | -60 | 39 | 781 | 4.13 |
| Right Superior Lateral frontal ** | 20 | 46 | 30 | 1118 | 4.03 |
| Distance < Natural |  |  |  |  |  |
| No brain region |  |  |  |  |  |

\*\* cluster reported at $p < 0.05$ FWE whole brain cluster corrected (initial cluster-forming threshold of $p < 0.001$ , uncorrected and minimum extent $k = 40$ )

Comparison of the Distance condition with the Natural condition (Distance > Natural) revealed a significant increase of the BOLD signal in the bilateral superior PFC (x,y,z: −24, 52, 22; 20, 46, 30; t = 6.76 and 4.94 for Left and Right respectively), the right dlPFC (x,y,z: 44, 16, 46; t = 5.35), the anterior cingulate cortex/supplementary motor area (ACC/SMA) (x,y,z: 3, 8, 62; t = 5.20) and bilateral angular gyrus (x,y,z: 56, −54, 27; −46, −60, 39; t = 4.22 and t = 4.13 for right and left respectively) (p<0.05 FWE whole brain cluster corrected) (Figure 4, Table 1). Again, we also plotted the beta parameters from these regions in the three different conditions (Figure 4). Finally, investigation of the contrast (Natural > Distance) revealed no brain region showing greater activity in Natural trials as compared to Distance trials.

**Figure 4:**
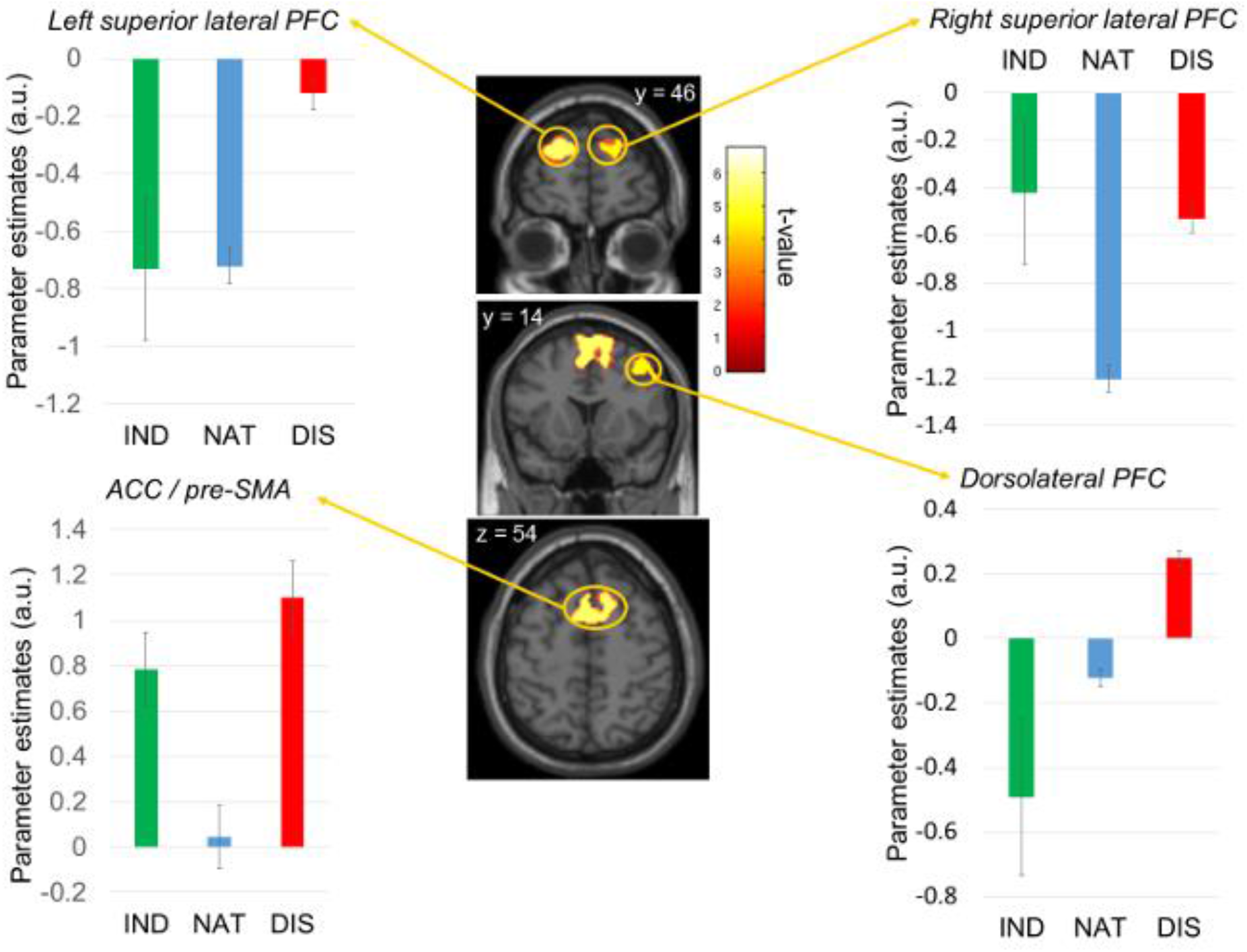
Down-regulation during the Distance condition increases activity of the bilateral superior PFC and the right dlPFC. From top to bottom: Left superior lateral PFC (x y z: −24 52 22) and right superior lateral PFC (x,y,z: 20, 46, 30); right dlPFC (x y z: 44 16 46) and ACC/pre-SMA (x,y,z: 3, 8, 62). Graphs indicate beta values extracted from clusters of activity. All activations are reported at a voxel-wise threshold of p<0.001 and cluster size > 40, which corresponded to a whole brain FWE cluster corrected threshold of p<0.05. Error bars show SEM.

##### Relationships between Ghrelin/leptin levels and brain activity related to different regulatory conditions

Because up-regulation during the Indulge condition selectively increased activity in the dorsomedial dmPFC, we thought to investigate the link between this brain response and leptin/ghrelin levels. To determine the potential relationship between hormone levels and regulatory mechanisms, we conducted a correlation analysis between the beta extracted from the dmPFC ROI (8 mm sphere centered on the peak dmPFC cluster identified in the comparisons Indulge>Natural) and leptin/grehlin levels. We observed a positive correlation between total ghrelin and betas extracted from the dmPFC ROI in the Indulge (r = 0.469, p = 0.024) and between ghrelin and betas in the Natural condition (r = 0.511, p = 0.013). No significant correlation was found between beta parameters from regions revealed by the contrast Distance > Natural and ghrelin level (Figure. 3B). The same procedure was conducted for leptin level but no significant correlations were revealed. Finally, we defined the ‘regulatory success of Willingness To Pay’ (WTP) as the WTP difference between the Indulge and Natural conditions for a given food (represented by an odor followed by a picture) in each case where participants bid more in the Indulge condition compared to Natural condition. The same procedure was used to compute the absolute value of the difference in WTP between Distance and Natural conditions for the cases where participants bid less in the Distance condition than in the Natural condition. We then conducted Pearson correlation analysis to determine if the ghrelin level is correlated with regulatory success when up- or down regulating. We observed a significant correlation between ghrelin levels and up-regulation (r= 0.373; p = 0.002) but not between ghrelin levels and down-regulation (p = 0,516) (Figure 3C).

##### Subjective value computation at the time of Willingness to Pay

Next, we searched for brain areas engaged in odor/image value computation using GLM2. We found that the vmPFC (x,y,z: 6, 42, 3; t = 3.96), and bilateral VSTR (x,y,z: −6, 14, −4; t = 4.20 and 6, 18, −2; t = 3.62 respectively left and right) positively correlated with participants’ bid in the Natural condition (p<0.05, FWE corrected within small volume correction) (Figure. 5, Table 2). Note that we used Natural trials only to determine brain areas correlating with WTP because there was not enough variation in the bids in the Indulge and Distance conditions to perform regression analyses between WTP and the BOLD signal. Indeed, WTP were always high in the Indulge condition and always low in the Distance condition, relative to the Natural condition. This prevented us to observe value signals across all conditions and to test for changes in slopes between value representation (as indexed by brain regressions with WTP) and regulatory conditions. Thus, we defined ROIs to investigate whether regulatory instructions modulated activity in the vmPFC and bilateral striatum. That is, we defined vmPFC and bilateral VSTR as spherical ROIs based on previous analyses reporting these regions as key areas for valuation (Clithero and Rangel, 2013; Hutcherson et al., 2012; Kober and Mell, 2015; Metereau and Dreher, 2015; Todd A. Hare, Colin F. Camerer, 2007) Within these ROIs, we used GLM1 to search for significant differences between regulatory instructions. The results revealed no significant differences across conditions within these ROIs. To illustrate this, we extracted beta parameters from these ROIs in the three different conditions and plotted them (Figure 5).

**Figure 5:**
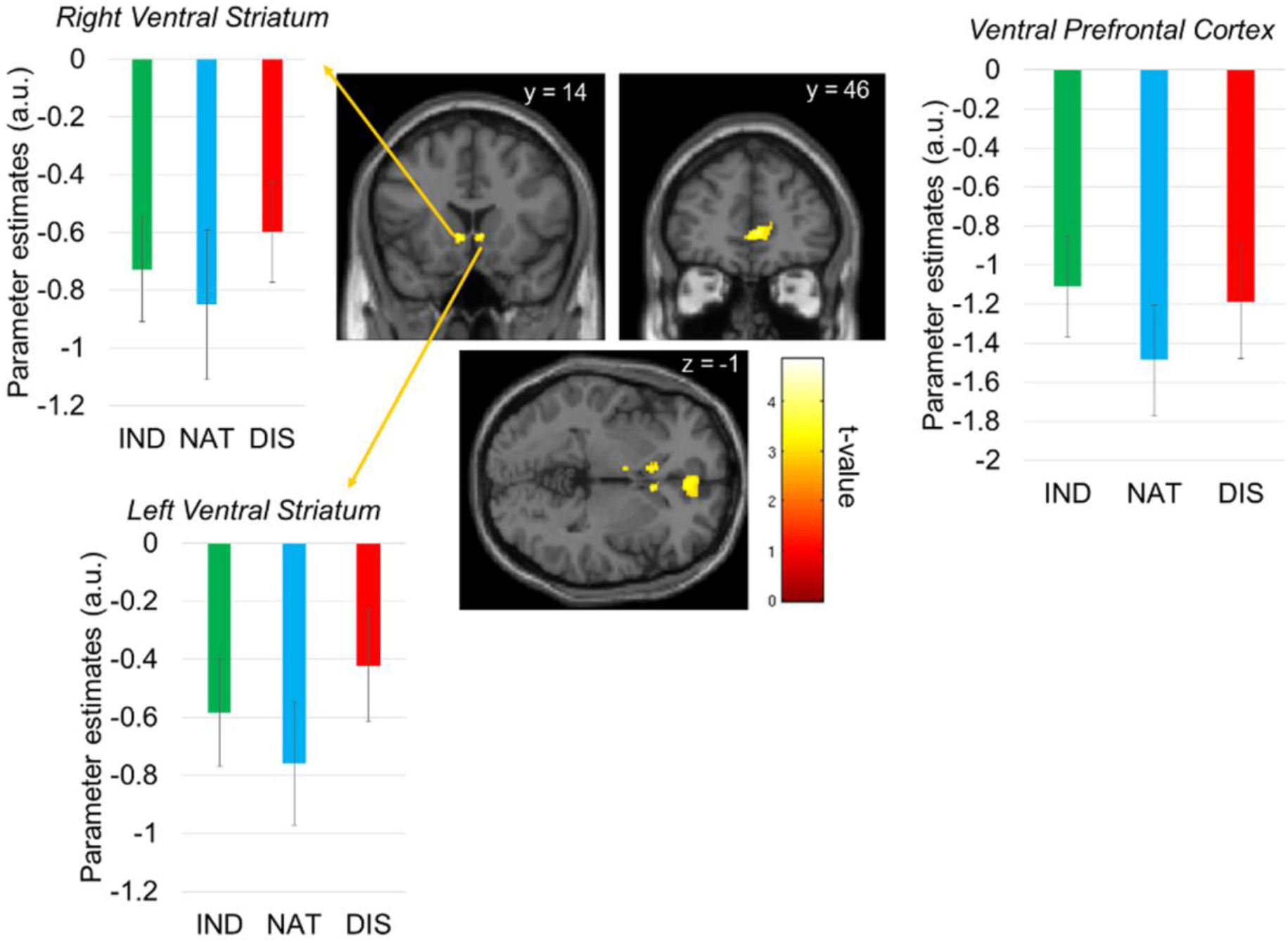
vmPFC and bilateral striatum correlate with bids in the Natural condition. The vmPFC (x,y,z: 6, 42, 3) and bilateral striatum (x,y,z: −6, 14, −4 and 6, 18, −2) correlate with the willingness to pay during the Natural condition with no differential effect of the regulation strategies on these regressions. All activations are reported at a whole brain FWE peak corrected threshold of p<0.05. Here the activation map is presented at p<0.001 uncorrected for display. Beta extracted within these regions come from GLM 2. Error bars show SEM.

**Table 2:** Brain areas correlating parametrically to the bid. ROI analyses were performed using a family wise error (FEW) peak cluster corrected for multiple comparisons. *small volume correction within a sphere ROI of 8 mm centered on peak activity from previous literature (Clithero and Rangel, 2013; Metereau and Dreher, 2015).

|  | MNI peak cluster coordinates |  |  |  |  |
| --- | --- | --- | --- | --- | --- |
|  | x | y | z | k | Z score |
| Ventromedial PFC * | 6 | 42 | 3 | 1229 | 3.42 |
| Left Ventral Striatum * | 6 | 18 | -2 | 627 | 3.19 |
| Right Ventral Striatum * | -6 | 14 | -4 | 619 | 3.58 |
| WTP x condition interaction | 6 | 42 | 3 |  |  |
| No brain region | 6 | 18 | -2 |  |  |

\* $p < 0.05$ , small-volume corrected within an ROI defined based on literature.

##### Neural representations of individual odor ranking

To test for brain regions involved in the individual ranking preferences of food categories, we used GLM3 (see methods) classifying food categories for each participant from the most preferred to the least preferred food category, based on mean bid, regardless of condition. Re-ranking each odor from the least preferred odor (R2) to the most preferred odor (R5) for each individual subject, we then tested for individual processing of food categories using linear combination: R2; R3; R4 and R5, and then used a regression test R2<R3<R4<R5 at the group level to identify brain regions encoding individual ranking preferences of food during the odor presentation. This analysis revealed that the right amygdala (x,y,z: 24, 2, −20; t =4.76), and bilateral occipital cortices (x,y,z: −24, −96, −12 / 22, −93, −9) correlated robustly and positively with food ranking preferences at the time of odor presentation (Figure 6, Table 3, p<0.05 FWE whole brain peak cluster corrected). The left amygdala was also found to be engaged in this correlation at a lower threshold (p<0.005, uncorrected).

**Figure 6:**
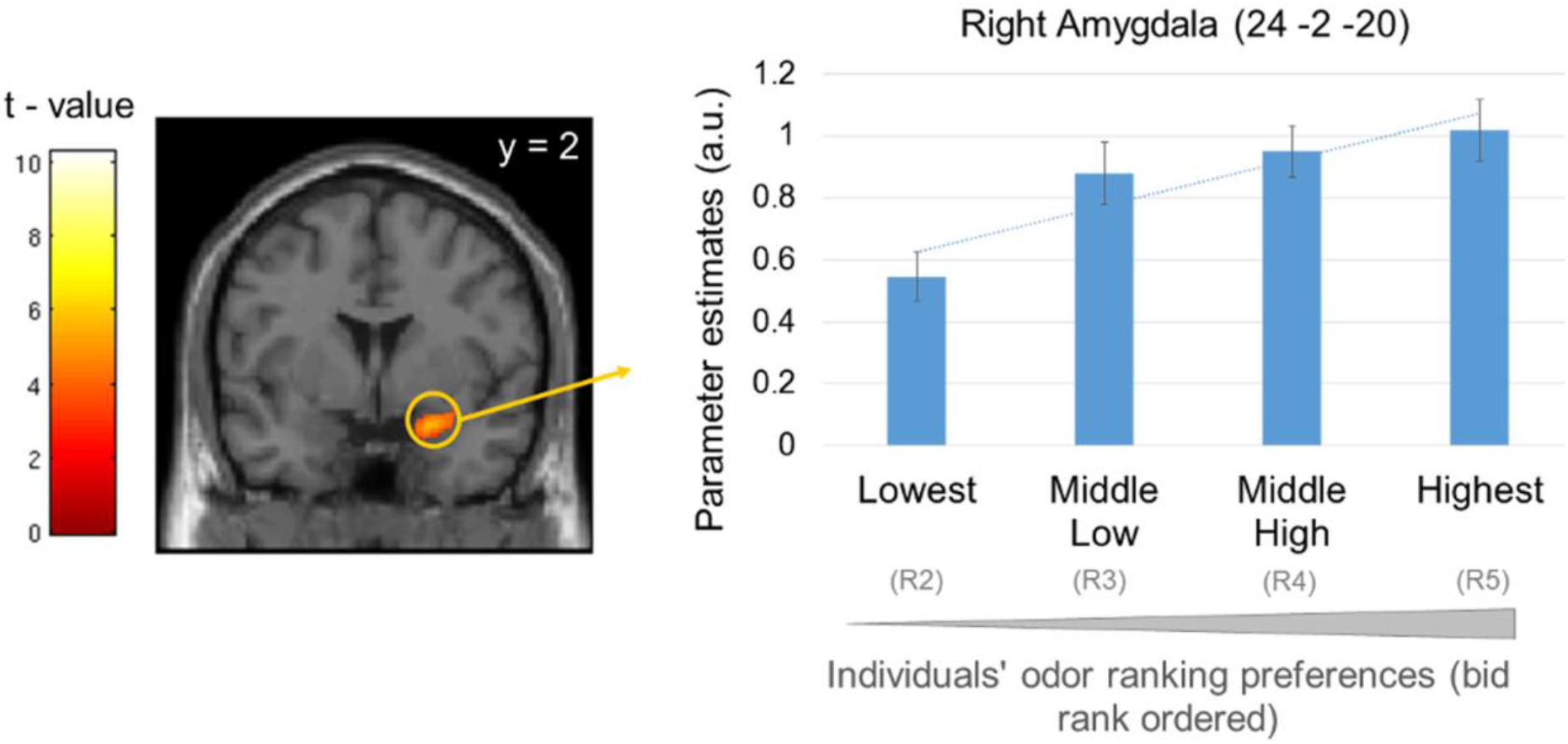
Amygdala activity correlates with individual food category ranking during odor presentation. *Left*. Right amygdala activity increased with higher individual ranking (x,y,z: 26, −2, −18). Activations are reported at a whole brain FWE peak corrected threshold of p<0.05. Similar correlation was observed in the left amygdala at a lower threshold (x,y,z: −21, 5, −18; p<0.005 uncorrected). Here the activation map is presented at p<0.005 unc for display purpose only. *Right*. Beta extracted in right the amygdala for each food category. Error bars show SEM.

**Table 3:** Brain regions correlating to food ranking preferences for each participant. **Cluster are reported at p < 0.05, family-wise error (FWE) cluster-level corrected for multiple comparisons (with an initial cluster-forming threshold of p < 0.001 and an extent k = 40 voxels). † Cluster are reported at p < 0.05, family-wise error (FWE) peak-level corrected for multiple comparisons (with an initial cluster-forming threshold of p < 0.001 and an extent k = 40 voxels).

|  | MNI peak cluster coordinates |  |  |  |  |
| --- | --- | --- | --- | --- | --- |
|  | x | y | z | k | Z score |
| During odor presentation |  |  |  |  |  |
| Right Amygdala † | 24 | -2 | -20 | 348 | 5.56 |
| Right Occipital ** | -24 | -96 | -12 | 4429 | 10.26 |
| Left Occipital ** | 22 | -93 | -9 | 3439 | 8.01 |
| During picture presentation |  |  |  |  |  |
| Right Occipital ** | -36 | -57 | -15 | 48576 | 23.42 |
| Middle Cingulate cortex ** | -6 | 14 | 45 | 3686 | 13.07 |
| Dorsolateral Prefrontal cortex ** | 38 | 6 | 36 | 2363 | 7.19 |
| Right Anterior Insula ** | 32 | 30 | 2 | 782 | 6.07 |
| Left Anterior Insula † | -30 | 26 | -4 | 562 | 5.92 |
| Right Inferior Parietal cortex † | 3 | -30 | -6 | 486 | 4.73 |
| Food preferences x condition |  |  |  |  |  |
| interaction (odor) |  |  |  |  |  |
| No brain region |  |  |  |  |  |
| Food preferences x condition |  |  |  |  |  |
| interaction (picture) |  |  |  |  |  |
| No brain region |  |  |  |  |  |
| During odor presentation |  |  |  |  |  |

\*\* cluster reported at $p < 0.05$ FWE whole brain cluster corrected (initial cluster-forming threshold of $p < 0.001$ , uncorrected and minimum extent $k = 40$ )
† peak activity reported at $p < 0.05$ FWE whole brain peak cluster corrected initial cluster-forming threshold of ( $p < 0.001$ , uncorrected and minimum extent $k = 40$ )

To test for any effect of conditions-by-preferences interaction we used a flexible factorial design including regressors denoting individual ranking preferences per condition during odor presentation. No significant BOLD response was observed in any brain areas supporting individual ranking preferences (Table 3, p<0.05 FWE whole brain cluster corrected).

##### Functional connectivity analysis

To test for changing functional connectivity according to regulation conditions between brain regions engaged in cognitive regulation (i.e. dlPFC and dmPFC identified with the GLM1) and a core component of the valuation network, i.e. the vmPFC, we performed a generalized PsychoPhysiological Interaction (gPPI) using the CONN toolbox. A gPPI model allows us to explore the physiological response (HRF convolved BOLD signal) in one region of the brain in terms of the context-dependent response of another region. This effectively provides a measure of task-modulated connectivity between two or more regions. Here, we conducted two ROI-to-ROI gPPI analysis using the CONN Toolbox (www.nitrc.org/projects/conn,RRID:SCR_009550). The first analysis was performed to investigate the functional connectivity between the dmPFC and the vmPFC during the Indulge compared to the Natural conditions, taking the dmPFC as a seed region (gPPI-1). The second analysis investigated the functional connectivity between the bilateral superior PFC, the dlPFC and the vmPFC during the Distance compared to the Natural conditions (gPPI-2).

The results revealed no significant differences in functional connectivity between the cognitive regulation brain regions identified using the GLM1 and the vmPFC during the perception of the combined olfacto-visual stimuli during the Indulge regulation compare to the Natural condition (Table 4). They neither revealed significant differences in functional connectivity between the right dlPFC or bilateral superior PFC and the vmPFC during the Distance condition compared to the Natural condition (peak voxel-level at p < 0.001, corrected at the cluster-level using a family-wise error rate (FDR) of p < 0.05 two-sided) (Table 4).

**Table 4:** Results from the ROI-to-ROI gPPI functional connectivity analysis. No significant functional connectivity was identified between regulatory regions and the vmPFC. Initial voxel-level at p < 0.005 for cluster formation, and then corrected at the cluster-level FDR<0.05.

|  | Seed | Target ROIs | p-val<br>unc. | p-val<br>FDR | t-value |
| --- | --- | --- | --- | --- | --- |
| gPPI - 1<br>[Indulge>Natural] | dmPFC | vmPFC | 0.84 | 0.96 | -0.2 |
| gPPI - 2 | Left IPFC | vmPFC | 0.83 | 0.83 | -0.21 |
| [Distance>Natural] | Right IPFC | vmPFC | 0.71 | 0.99 | -0.38 |
|  | dRight dIPFC | vmPFC | 0.83 | 0.83 | -0.21 |

## Discussion

One important aspect of this study was to investigate, in food deprived participants, the neural mechanisms engaged in cognitive regulation of food stimuli presented in the visual and olfactory domains. Both willingness to pay for food and sniffing parameters were modulated by regulatory conditions, indicating reliable regulatory mechanisms at the behavioral and physiological levels. At the brain system level, increased dorsomedial PFC activity occurred during up-regulation of appetite (Figure 3A), whereas a brain network, including the bilateral lPFC, the right dlPFC and the ACC was more engaged during down-regulation of appetite as compared to the Natural condition (Figure 4). Activity from to the valuation system, including the vmPFC and bilateral striatum correlated with increasing WTP assigned to the food, but was not modulated by appetite regulatory instructions, confirming that engagement of this brain system is relatively automatic (Figure 5). Finally, the amygdala response positively correlated with the individual preference for the specific food presented, and this ‘odor-specific preference’ response was also resilient to neuroregulatory instructions. Together, these results demonstrate the existence of separate brain systems responding or not to appetite regulation when subjects are hungry.

Our findings provide novel insights into the neurobiological mechanisms involved in the cognitive regulation of ecological bimodal food cues. First, cognitive regulation modulated the duration (Figure 2D) and the amplitude (Figure 2E) of the first inspiration, showing that physiological parameters of the olfactory system are under modulation of cognitive regulation. Our results extend early findings on cognitive regulatory mechanisms, and highlight the fact that human sniffing is influenced by internal states, such as motivation or homeostatic state. For example, hunger increases sniff duration compared to satiety, even when sniffing clean air (Prescott et al., 2010). Cognitive regulation also modulated parameters of the entire sniffing period (Figures 2H, 2G). These results suggest that cognitive regulation can control sniff parameters and modulate behavior. This is consistent with the fact that breathing phase modulates discrimination of fearful faces, as fearful faces were recognized more quickly during expiration compared to inspiration (Zelano et al., 2016). Together, these inhalation results suggest that subjects have meta-cognition about the impact of odors in their self-control, and that they try to influence their regulatory abilities by accessing this mechanism. This is an important finding because much remains to be learnt about the channels through which individuals exercise dietary control. Finally, participants’ WTP for food was higher in the Indulge condition, as compared to the Natural condition, whereas they bid less under the Distance condition (Figure 2A). These findings extend previous results restricted to the visual modality to multimodal food stimuli (Boswell et al., 2018; Hutcherson et al., 2012) and further show that humans are able to regulate their WTP when the food is presented in both olfactory and visual modalities.

The increased engagement of the dorsomedial PFC (dmPFC) observed with up-regulation of appetite towards food stimuli is consistent with the fact that this brain region shows a specific heightened response to food when subjects are hungry (Anderson et al., 2006; Giannopoulou et al., 2018; LaBar et al., 2001) (Figure 3A). This finding supports the idea that this brain region modulates motivation towards food. These results suggest that during up-regulation, activity within the dmPFC increases together with a concurrent increase in appetite towards food stimuli. In obese populations, a meta-analysis revealed higher activity of the dmPFC when viewing food pictures (Brooks et al., 2013) while patients suffering from Binge Eating Disorders rate food stimuli as significantly less desirable than healthy controls (Uher et al., 2004).

Conversely, the dlPFC, bilateral superior PFC and the ACC were more active when participants down-regulated their appetite towards food odor/image stimuli (Figure 4). The dlPFC is known to be engaged in regulation of appetite for visually presented food stimuli (Hutcherson et al., 2012a; Kober et al., 2010; Lewis and Bates, 2014, Tusche and Hutcherson, 2018) and the ACC plays a critical role in response inhibition and in the selection of appropriate behavior to resolve situations such as action suppression (Cole and Schneider, 2007; Simmonds et al., 2008). Moreover, the bilateral superior PFC has been associated with cognitive strategies to suppress the desire for food stimuli (Siep et al., 2012b). Engagement of a brain network including the dlPFC has been reported in emotional down-regulation of odors (Billot et al., 2017). Moreover, regulation-related neural activation patterns in a right dlPFC area has been shown to reliably predict how well participants decrease taste weights attribute in food choices (Tusche et al., 2018). Together, these findings are consistent with the hypothesis that cognitive regulation of appetite and emotional regulation may share common neural substrates because increased dlPFC response has been observed during down-regulation of negative emotion, and decreased dlPFC activity has been found during down-regulation of positive emotion (Ochsner and Gross, 2005).

When investigating the brain systems engaged in the valuation of food items in response to ecological cues in the absence of cognitive regulation (Natural condition), we observed engagement of the valuation brain system, consisting of the vmPFC and the bilateral striatum (Figure 5). These brain regions have previously been shown to be key for valuation of food items presented visually (Clithero and Rangel, 2013). We extend these previous results to multimodal situations in which food is experienced in the visual and olfactory modalities. Such vmPFC engagement in the valuation of a food item is much closer to experiencing real food (combining vision, smell and taste). This brain region has also been observed during anticipation of salient food (liquid) reinforcers that are really delivered inside the scanner (Metereau and Dreher, 2015; O’Doherty et al., 2002). Thus, vision and smell play a key role in constructing a unified and co-occurring percept defined as flavor when anticipating and experiencing food items.

It should be noted that none of the brain valuation regions were modulated by cognitive regulation. In fact, distinct brain regions were engaged during the Indulge and Distance conditions. This confirms that valuation is relatively automatic, as previously suggested (Lebreton et al., 2009) and that cognitive regulation of food odor and image does not modulate the valuation system itself. A number of previous studies using similar paradigms but only with food presented in the visual domain (Hare et al., 2009, 2011; Hutcherson et al., 2012; Krishna, 2012), have reported that regulation involves the modulation of value signals in the vmPFC, as well as interaction with regions like the dlPFC. To investigate whether this was the case in the current dataset, we performed a functional connectivity analysis between the regions engaged in cognitive regulation i and the vmPFC. Using ROI-to-ROI gPPI analysis, we investigated changes in connectivity pattern between the dmPFC and the vmPFC during the Indulge as compared to the Natural conditions. We also investigated changes in connectivity patterns between the right dlPFC, the bilateral superior lPFC and the vmPFC during the Distance as compared to the Natural conditions. These analyses failed to reveal that cognitive regulation is exerted by changed in functional connectivity between the dmPFC and the vmPFC during the Indulge vs Natural conditions. No functional change in connectivity was observed between the dlPFC or bilateral superior lPFC and vmPFC for the Distance vs Natural conditions either (see results section and Table 4). Although these findings may be surprising at first stake, a number of previous failures to observe changes in modulation of the vmPFC during cognitive regulation of decision making suggest an alternative hypothesis (Hollmann et al., 2012; Yokum and Stice, 2013; Tusche et al., 2018). It is possible that in our study, as in these previous studies, cognitive regulation alters value representations at a relatively low level, by amplifying or diminishing attribute representations directly in a distributed set of specific, dedicated attribute-coding areas. Consistent with this possibility, a recent study observed that cognitive regulation did not operate at higher levels in centralized, domain-general value integration area such as the vmPFC (Tusche and Hutcherson, 2018). Instead, cognitive regulation of decision-making altered value representations at a relatively low level, representing food attributes in a dlPFC region. Further work will be needed to understand the respective roles of the vmPFC and dlPFC, as well as their interactions during cognitive regulation of food stimuli presented in multimodal domains.

Investigation of the brain regions encoding individual odor preferences, based on rank ordering of the foods presented, showed that activity of the amygdala robustly increased as a function of increasing odor preference (Figure 6). The cognitive regulation conditions did not modulate this amygdala response, and there was also no interaction between regulatory instruction and odor ranking at the behavioral level. This amygdala response may thus correspond to a relatively automatic route that integrates information about potential outcome value and action-outcome association to guide choice behavior (Balleine and Killcross, 2006; O’Doherty, 2004; Saez et al., 2017; Seymour and Dolan, 2008). Consistent with this interpretation, a previous study showed that the amygdala encodes subjective valence of odors (Jin et al., 2015).

Finally, correlational analysis revealed a positive relationship between blood levels of ghrelin and BOLD response in the dmPFC that up-regulated the subjective value of food items in the Indulge condition (Figure 3B). Individuals with higher blood levels of ghrelin were better at exercising up-regulation as they showed higher up regulatory success when comparing bids from the Indulge vs Natural conditions (Figure 3C). Thus, the relationship between dmPFC activity and ghrelin levels may be a neurobiological marker for up-regulation success. Ghrelin regulates food intake (Date et al., 2001; Müller et al., 2015) and levels of ghrelin positively correlate with increased hunger and with increasing activity in large brain networks involved in the regulation of feeding and in the appetitive response to food cues (Batterham et al., 2007; Goldstone et al., 2004; Jones et al., 2012; Wei et al., 2015; Zanchi et al., 2017). All these studies showed increased neural response to food pictures in regions of the brain engaged in encoding the automatic incentive value of food cues, but did not investigate how ghrelin modulates brain regions engaged with up-regulation of food cues, as in the current study. Ghrelin may act on the brain through several mechanisms, including ghrelin receptors in the gut relaying information *via* the vagus nerve (Date, 2013), the hypothalamus regulating feeding behavior and the dopaminergic system (Abizaid et al., 2006; Perello et al, 2012). Our results suggest that the dmPFC plays a crucial role in the relationship between ghrelin and up-regulation of feeding behavior.

Together, our results demonstrate that in the context of hunger, up- and down-regulation of appetite towards realistic food stimuli presented in the olfactory and visual domains are mediated by distinct brain networks. The medial prefrontal cortex is engaged in up-regulation whereas the lateral prefrontal cortex is engaged in down-regulation. Our findings also provide new insights to the relationship between higher-level brain regions engaged in up-regulating food consumption and ghrelin.

## Materials and Methods

### Participants

Twenty five healthy volunteers (12 females, 13 males; age range 18-33 years; and mean age (M) 22.45 ± (SEM) 3.88) were recruited through a mailing list from the University of Claude Bernard Lyon 1. All participants had a normal Body Mass Index (BMI) (mean (M) 21.74 ± (SEM) 0.36). For inclusion in the study, participants were required to follow the following criteria: french-speaking, right-handed, no current medical treatment, no history of neurological or psychiatric disorders and no auditory, olfactory or visual deficits. Furthermore, volunteers were screened for general MRI contra-indications. A physician conducted medical examinations concerning inclusion criteria such as physical and psychological health. Participants gave their written consent and received monetary compensation for the completion of the study. This study was approved by the Medical Ethics Committee (CPP Sud-Est III, ID RCD: 2014-A011661-46).

### Stimuli and delivery

Olfactory stimuli (apricot, pineapple, dark chocolate, milk chocolate, all EURACLI products, Chasse-sur-Rhone, France; Respective concentration vol/vol: 75%, 25%, 75%, 75%) and corresponding visual stimuli (depicting desserts; 8 different images *per* odor type (Figure 1B) were presented using a device adapted for fMRI olfactory/visual experiments and described in details in Sezille et al. (Sezille et al., 2013). Airflow control, odor concentration and stimulus duration, as well as collection of participants’ responses were all managed by the system, which is composed of a series of modules: 1/ an airflow source, 2/ a diffusion module controlling odorant duration and concentration through regulation of airflow, 3/ a mixing head used to *(i)* mix non-odorized air from the first module with a specific odorant (controlled by the second module) and (ii) send the diluted odor to the nose, 4/ a software enabling presentation of verbal material (instructions) and visual stimuli, 5/ a response box to provide subjective ratings.

To ensure synchronization between fMRI measures and odor diffusion, olfactory stimuli were diffused at the beginning of each nasal inspiration. To this end, the respiratory signal was acquired using an airflow sensor that was integrated with an amplifier interface. A microbridge mass airflow (AWM2100V, Honeywell, MN, USA) allowed acquisition of both inhalation and exhalation phases. The airflow sensor was connected to a nasal cannula (Cardinal Health, OH, USA; 2.8 mm inner diameter tube) positioned in both nostrils. Sniffing was digitally recorded at 100 Hz and stored in the odor diffusion computer. Sniffs were pre-processed by removing baseline offsets, and aligned in time by setting the point when the sniff entered the inspiratory phase as time zero. Inspired volume, max amplitude rate and sniff durations were calculated for the first sniff of every trial. Mean sniffing parameters during the entire odor presentation was also recorded.

The whole system was controlled using LabVIEW^®^ software. A multiple function board (National Instruments, TX, USA) was used to acquire all experimental events (olfactory, visual, instructions), signals from the respiratory sensor and the response box, which allowed synchronization with the external system (fMRI scanner).

### Experimental design

Each trial started with a visual instruction indicating the type of trial (i.e., Indulge, Distance or Natural) for 3 s (seconds) (Figure 1A), followed by the diffusion of one of the four categories of odors (apricot, pineapple, dark chocolate or milk chocolate) for 3.2 ± 0.8s, and synchronized with the respiration of the participants. Afterwards, a visual image corresponding to the odor was presented for 2 s (e.g. a visual picture of a pineapple pie following the smell of pineapple) (Figure 1B). Finally, the participants had 5 s to select the price they were willing to pay for the presented food. Participants were able to choose one price from the five depicted on the screen (i.e. € 0.50, € 1.00, € 1.50 €2.00 or € 2.50); consistent with auction rules described by Becker-DeGroot-Marschack (BDM) (M. Becker, Morris H. DeGroot, 1964; Plassmann et al., 2007). Following participant’s response, a fixation cross was presented for 5.4 ± 0.6s.

Before the “Modulatory instruction and bidding task” began, participants received specific modulatory instructions for each trial type. For the Indulge condition, participants were asked to smell the odor and to keep looking at the presented food image while adopting thoughts that would increase their desire to eat the presented food immediately. For the Distance condition, participants were asked to smell the odor and keep looking at the presented food image while adopting thoughts that would decrease their desire to eat the presented food immediately. For the Natural condition, participants were asked to smell the odor and keep looking at the presented food image while allowing any thoughts and feelings that came naturally in that moment.

Before scanning, participants were asked to rate how hungry they felt on a continuous scale ranging from 0= “not hungry at all”, 50 = “moderately hungry”, to 100= “never been so hungry”. They also rated the quantity of food they were able to eat, based on a similar continuous scale range from 0, “I cannot eat anything”, to 100, “I could eat anything”. This allowed us to check the subjective hunger level of each participant. The fMRI task consisted of 120 trials divided into four sessions. Each session included 30 trials, 24 trials divided into the three trial types (i.e. 8 trials for each condition) and six resting trials called Air-clean (i.e. in which no instruction, no odor and no image were presented). Each session comprised a fixed order of presentation of 30 trials. The four sessions were presented randomly to participants.

After scanning, blood samples were drawn by a nurse to latter assess ghrelin and Leptin levels.

### fMRI data acquisition

All MRI acquisitions were performed on a 3 Tesla scanner using EPI BOLD sequences and T1 sequences at high resolution. Scans were performed on a Siemens Magnetom Prisma scanner HealthCare, CERMEP Bron (single-shot EPI, TR / TE = 2500/21, flip angle 80 °, 45 axial slices interlaced 2 mm thickness 2 mm gap, FOV = 232 mm and 116 die). A total of 1120 volumes were collected over four sessions during the experiment, in an interleaved ascending manner. The first acquisition was done after stabilization of the signal. Whole-brain high-resolution T1-weighted structural scans (1 x 1 x 1 mm) were acquired for each subject, co-registered with their mean EPI images and averaged across subjects to permit anatomical localization of functional activations at the group level. Field map scans were acquired to obtain magnetization values that were used to correct for field inhomogeneity.

### fMRI data preprocessing

Image analysis was performed using SPM12 (Wellcome Department of Imaging Neuroscience, Institute of Neurology, London, UK, fil.ion.ucl.ac.uk/spm/software/spm12/). Time-series images were registered in a 3D space to minimize any effect that could result from participant head-motion. Once DICOMs were imported, functional scans were realigned to the first volume, corrected for slice timing and unwarped to correct for geometric distortions. Inhomogeneous distortions-related correction maps were created using the phase of non-EPI gradient echo images measured at two echo times (5.40 ms for the first echo and 7.86 ms for the second). Finally, in order to perform group and individual comparisons, they were co-registered with structural maps and spatially normalized into the standard Montreal Neurological Institute (MNI) atlas space. Following this, images were spatially smoothed with an 8 mm isotropic full-width at half-maximum (FWHM) Gaussian kernel using standard procedures in SPM12.

### fMRI data analysis and imaging statistics

To address the questions raised in the introduction, that is whether: (1) cognitive regulation modulates both sniffing and bidding behavior; 2) a common valuation system is involved in the valuation of odor/image stimuli; 3) there are distinct brain regions (especially prefrontal regions: dmPFC vs dlPFC) supporting up- and down-regulation), we estimated three general linear models (GLMs). Each GLM was estimated in three steps. First, we estimated the model separately for each individual. Second, we calculated contrast statistics at the individual level. Third, we computed second-level statistics by carrying out various statistical tests on the single-subject contrast coefficients.

Statistical analyses were performed using a conventional two-level random-effects approach with SPM12. All GLMs included the six motion parameters estimated from the realignment step. Statistical inference was performed at a standard threshold of p < 0.05, family-wise error (FWE) cluster-level corrected for multiple comparisons, with an initial cluster-forming threshold of p < 0.001 and an extent k = 40 voxels.

#### Analysis of cognitive regulation during odor/image presentation

To determine the brain regions involved in cognitive regulation during the odor/image presentation, we used GLM1. This GLM1 consisted of 9 regressors of interest. Regressors R1–R3 modeled brain response related to the instructions according to the condition, respectively Natural (R1), Indulge (R2) and Distance (R3). R1-R3 were modeled as a boxcar function time-locked to the onset of the instruction with duration of 3 s. R4 to R6 denoted regressors during food stimuli delivery in Natural (R4), Indulge (R5) and Distance (R6) trials and were modeled as a boxcar function beginning at odor presentation and during the entire period of food odor/image presentation (mean 8.4 ± 1.6s). Finally, R7 to R9 modeled brain response related to the rating in the three regulatory instructions Natural (R7), Indulge (R8) and Distance (R9). R7-R9 were modeled as a boxcar function time-locked to the onset of the rating (willingness to pay) period with duration of response times (RTs: 1.8 ± 0.5s). Missed trials were modeled as a separate regressor over the duration of the entire trial. Finally Air-clean trials were modeled separately using three distinct regressors. R10, denoting the instructions period from the Air-clean trials and modeled as a boxcar function time locked at the beginning of the instructions and during 3 seconds. R11, that denoted the stimulus period from the Air-clean trials, starting from the beginning of stimulus (even if the stimulus is a blank odor followed by a dark screen) and during 8 seconds. And R12, that denoted the rating period from the Air-clean trials and modeled as a stick function (because participants didn’t have to indicate their willingness to pay). The model also included motion parameters and session constants as regressors of no interest. To test for cognitive regulation, we computed the following contrasts: [Indulge (R5) > Natural (R4)] (Figure 3; *Table 1*), [Distance (R6) > Natural (R4)] (Figure. 4; Table 1) at the single level and then used a one-sample t-test at the group level on the single-subject contrast coefficients estimated. We also computed the opposite contrasts at the first level [Natural (R4) > Indulge (R5)] and [Natural (R4) > Distance (R6)] and then similarly used a one-sample t-test at the group level on the single-subject contrast coefficients.

#### Analysis of value computation in the Natural context

We used GLM2 in order to investigate brain regions involved in the computation of value while experiencing food odor and image stimuli. We investigated the brain regions reflecting such value computation by searching for brain areas in which the BOLD response correlated with bids during the Natural trials. GLM2 had one regressor of interest R1, consisting of the values of participants’ bids in the Natural trials. The hemodynamic response of this categorical function was convolved with a boxcar beginning at the time of the first odor inspiration and terminating at the bid response (average duration of 10,2 ± 1.75s). The instruction period was regressed using a boxcar function, starting from the beginning of instructions with a duration of 3s. GLM2 also includes regressors denoting others conditions stimuli (i.e. Indulge and Distance). This regressor consisted of a boxcar function starting from the beginning of inspiration and lasting until the end of rating (average duration of 10.2 ± 1.78s). The model also included motion parameters and session constants as regressors of no interest. Finally, Air-clean and missed trials were modeled separately with a duration lasting for the entire trial. In order to reveal brain areas involved in the computation of value, contrasts on the bid parametric modulator on Natural trials were computed. Then, a one-sample t-test was performed at the group level on single-subject contrast coefficients (Figure 5; Table 2).

We had strong *a priori* interest concerning the vmPFC and the ventral striatum because previous studies revealed that these regions perform value computation (Hare et al., 2009; Hutcherson et al., 2012; Kober and Mell, 2015; Metereau and Dreher, 2015). Region of Interest (ROI) analysis was thus conducted in a vmPFC ROI defined as an 8-mm diameter sphere, centered at x,y,z = −2, 40, 2, based on a previous meta-analysis study showing that this region is involved in the processing of food value presented visually (Clithero and Rangel, 2013), leading to a vmPFC ROI of 573 voxels. Based on this same study, we also defined two ventral striatum ROIs (left VSTR, defined as an 8-mm diameter sphere, centered at x,y,z = −8, 8, −6; right VSTR defined as an 8-mm diameter sphere, centered at x,y,z = 10, 14, −4, both including 573 voxels. All ROI were defined using WFU_PickAtlas (http://fmri.wfubmc.edu/software/PickAtlas). After ROIs creation, they were coregister on the functional images in order to keep voxel size.

#### Analysis of odor preferences encoding

To test for brain regions involved in the subjective preferences of food categories during odor perception, we defined a last GLM, designated GLM3. First of all, because there is no interaction between food category bidding behavior and regulatory conditions, we classified food stimuli categories (i.e. apricot, pineapple, milk chocolate, dark chocolate) from the least preferred to the most preferred, based on the mean bid regardless of the condition for all subjects. This allowed us to classify food categories for twenty-one participants. We were not able to classify food categories for 3 participants because of equal average WTP for at least 2 food categories. This GLM3 is composed of one regressor R1 denoting the instructions period, consisting of a boxcar function starting at the beginning of a trial and lasting 3 s. GLM3 also included 4 regressors of interest R2-R5 denoting the least preferred food odor categories (R2), the second less preferred food odor categories (R3), the second most preferred food odor categories (R4) and the most preferred food odor categories (R5). The hemodynamic response was convolved with a boxcar function beginning at the first inspiration at the odor presentation and terminating at the end of the odor presentation(duration 3.2 ± 0.8s). GLM3 also includes regressors denoting food categories presented visually and ranked similarly to the odor (R6 to R9). The hemodynamic response was convolved with a boxcar function beginning at the picture presentation and lasting for 2 seconds duration. Additionally, the rating period was modeled with a boxcar function starting from the beginning of the rating period and during the entire rating period (1.8 ± 0.61). The model also included motion parameters and session constants as regressors of no interest. To test for brain regions processing individual ranking preferences of food odor categories, we computed the following linear combination: R2, R3, R4 and R5 at the individual level. Then, we used a regression test R2<R3<R4<R5 at the group level analysis to identify brain regions encoding individual ranking preferences of odors (Figure 6; Table 3).

#### Functional connectivity between valuation and regulatory regions

A classical study reported that the dlPFC indirectly modulates the vmPFC via the iFG to exert self-control during food choices (Hare et al., 2011). However, a number of subsequent studies did not observe modulation of the vmPFC during cognitive regulation by regulatory regions (Hollmann et al., 2012; Yokum and Stice, 2013; Tusche et al., 2018). Here, we tested whether brain regions involved in the valuation of food cues are modulated by regulatory instructions during stimuli perception. To do so, we conducted two ROI-to-ROI generalized Psychophysiological Interaction (gPPI), between regulatory regions (i.e. dmPFC for the Indulge condition and bilateral superior lPFC and right dlPFC for the Distance condition) and valuation regions (vmPFC and bilateral striatum) using the CONN Toolbox (www.nitrc.org/projects/conn,RRID:SCR_009550). This allowed us to explore the changes in connectivity patterns between the regulation and valuation regions according to the regulatory instructions and so, to determine if valuation regions were differentially connected to regulation regions according to the regulatory goals demand. To perform this analysis, preprocessed functional images obtained from SPM as well as the design matrix coming from the GLM1 were loaded into the CONN toolbox. The CONN toolbox implemented the anatomical component-based noise correction method (Behzadi, 2008), extracting principal components related to the segmented CSF and white matter. This approach has been shown to increase the validity, sensitivity and specificity of functional connectivity analyses (Chai et al., 2012). Therefore, white matter and CSF noise components as well as motion parameters (six dimensions with first temporal derivatives resulting in twelve parameters) were regressed out during the denoising step. To control for simple condition-related activation effects, we also included the main task effects (related to GLM1 regressors) as confound regressors during the denoising step.

Two gPPI analyses were performed to investigate, in one hand, the difference in functional connectivity between the vmPFC and dmPFC identified with the contrast [Indulge (R5) > Natural (R4)] and, on another hand, the difference in functional connectivity between the vmPFC and the regulation regions identified in the contrast [Distance (R6) > Natural (R4)] with the GLM1. Concerning the Indulge regulation condition, we defined an 8 mm sphere ROI centered on the peak activity (x,y,z:- 4,58,21) from the one-sample t-test [Indulge (R5) > Natural (R4)] contrast. For the valuation region, we used the ROI of the vmPFC used for the GLM2 and entered it as target for the first gPPI ROI-to-ROI analysis gPPI-1 (Table 4).

For the Distance regulation condition, three others 8-mm diameter sphere ROIs were constructed from the clusters identified in the one-sample t-test [Distance (R6) > Natural (R4)] contrast, centered on the peak activity from the right lPFC, left lPFC and dlPFC (respectively x,y,z: 20,46,30; −24,52,22 and 44,16,46). Here again, the vmPFC ROI from the GLM2 was used as target in this second gPPI ROI-to-ROI analysis gPPI-2 (Table 4).

To account for false positives in multiple comparisons, results were thresholded at the peak voxel-level at p < 0.001, and then corrected at the cluster-level using a family-wise error rate (FDR) of p < 0.05 two-sided.

### Behavioral analysis

Due to excessive head motion, one participant was removed from the fMRI analyses (resulting in n=24 for fMRI analysis). Another participant had to be excluded from the leptin/ghrelin Pearson correlation analysis because the hormonal assessment data from this participant was missing (resulting in n=23 for this analysis).

All statistical analyses were performed using SPSS v21.0 (SPSS Inc., Chicago, IL, USA). Normal distribution was assessed with a Shapiro-Wilk test. If data distribution was not normal, we performed a Friedman test, otherwise a repeated measure ANOVA was conducted. Then, we ensured that homoscedasticity of variances were respected using a Mauchly test. If not, we applied a Greenhouse-Geisser correction to our ANOVA. For multiple comparisons, *post-hoc* (with Bonferroni correction) comparison was conducted according to the previous test used.

## Acknowledgments

This work was performed within the framework of the Laboratory of Excellence LABEX ANR-11-LABEX-0042 of Université de Lyon, within the program “Investissement d’Avenir” (ANR-11-IDEX-0007) operated by the French National Research Agency (ANR). This work was also supported by grants from the Agence Nationale pour la Recherche (ANR Social-POMDP). We thank the staff of CERMEP–Imagerie du Vivant (Lyon) for helpful assistance with data collection.

## Author Contributions

Conceptualization, J.C.D., M. B.; Methodology, J.C.D, R. J., M. F.; Investigation, M.F., A. F, M.B.; Resources, J.C.D and M.B.; Writing – Original draft, R.J.; M. B. and J.C.D.; Writing – Review & Editing, R.J; E.D. and J.C.D.; Funding Acquisition, J.C.D.; Supervision, J.C.D.

